# Neuromodulation enables temperature robustness and coupling between fast and slow oscillator circuits in *Cancer borealis*

**DOI:** 10.1101/2021.11.04.467352

**Authors:** Carola Städele, Wolfgang Stein

## Abstract

Acute temperature changes can disrupt neuronal activity and coordination with severe consequences for animal behavior and survival. Nonetheless, two rhythmic neuronal circuits in the crustacean stomatogastric ganglion (STG) and their coordination are maintained across a broad temperature range. However, it remains unclear how this temperature robustness is achieved. Here, we dissociate temperature effects on the rhythm generating circuits from those of upstream ganglia. We demonstrate that heat-activated factors extrinsic to the rhythm generators are essential to the slow gastric mill rhythm’s temperature robustness and contribute to the temperature response of the fast pyloric rhythm. The gastric mill rhythm crashed when only the STG circuits were heated. It could be restored when upstream ganglia were heated in addition, and the activity of the peptidergic modulatory projection neuron (MCN1) increased. Correspondingly, MCN1’s neuropeptide transmitter stabilized the rhythm and maintained it over a broad temperature range. Extrinsic neuromodulation is thus essential for the oscillatory circuits in the STG and enables neural circuits to maintain function in temperature-compromised conditions. In contrast, integer coupling between pyloric and gastric mill rhythms was independent of whether extrinsic inputs and STG pattern generators were temperature-matched or not, demonstrating that the temperature robustness of the coupling is enabled by properties intrinsic to the rhythm generators. However, at near-crash temperature, integer coupling was maintained only in some animals but was absent in others. This was true despite regular rhythmic activity in all animals, supporting that degenerate circuit properties result in idiosyncratic responses to environmental challenges.

## Introduction

Acute temperature changes are a significant challenge to neurons and neuronal circuits because they alter active and passive biophysical membrane properties but rarely linearly or equally across properties *(Cao and Oertel, 2005; Collins and Rojas, 1982; Hille, 1978)*. Consequently, temperature changes can disrupt neuronal activity with severe consequences for behavior and survival. All levels of neuronal processing are affected by temperature changes, including synaptic release, coupling, and dendritic and somatic integration. Possessing compensatory mechanisms that protect neurons from detrimental temperature changes are thus vital components of survival, particularly for poikilothermic animals, which can experience significant and rapid environmental temperature fluctuations.

*Powell et al. (2021)* have recently demonstrated that the coordination between two oscillatory circuits in the stomatogastric nervous system (STNS, Fig. 1A) can be maintained across a broad range of temperatures (7 to 23°C). Coupled oscillators are ubiquitous in nervous systems, and coordination between them is often necessary for proper function. In the STNS of the Jonah crab *Cancer borealis*, the gastric mill and pyloric central pattern generators (CPGs) are coupled despite these rhythms running at very different speeds *(Bartos et al., 1999; Nadim et al., 1998; Stein, 2017)*. The episodic slower gastric mill rhythm has a cycle period of ∼10s, while the continuous pyloric rhythm is about ten times faster (∼1s). Reciprocal inhibition of the lateral gastric (LG) neuron and its half-center antagonist Interneuron 1 (Int1) generates the gastric mill rhythm. The pyloric rhythm is generated by pacemaker neurons, including the anterior burster (AB) and two pyloric dilator (PD) neurons. However, the gastric mill rhythm requires neuromodulatory input from descending projection neurons to be activated *(Stein, 2009)*. When both rhythms are active, the gastric mill cycle period is an integer multiple of the pyloric cycle period (=integer coupled). This integer coupling is mainly mediated by the pyloric pacemakers’ synaptic inhibition to Int1. Rhythmic pacemaker activity inhibits Int1 (Fig. 1B), which results in disinhibition of LG, enabling LG to burst in synchrony with the pyloric pacemakers *(Bartos et al., 1999; Nadim et al., 1998)*.

**Figure 1.**
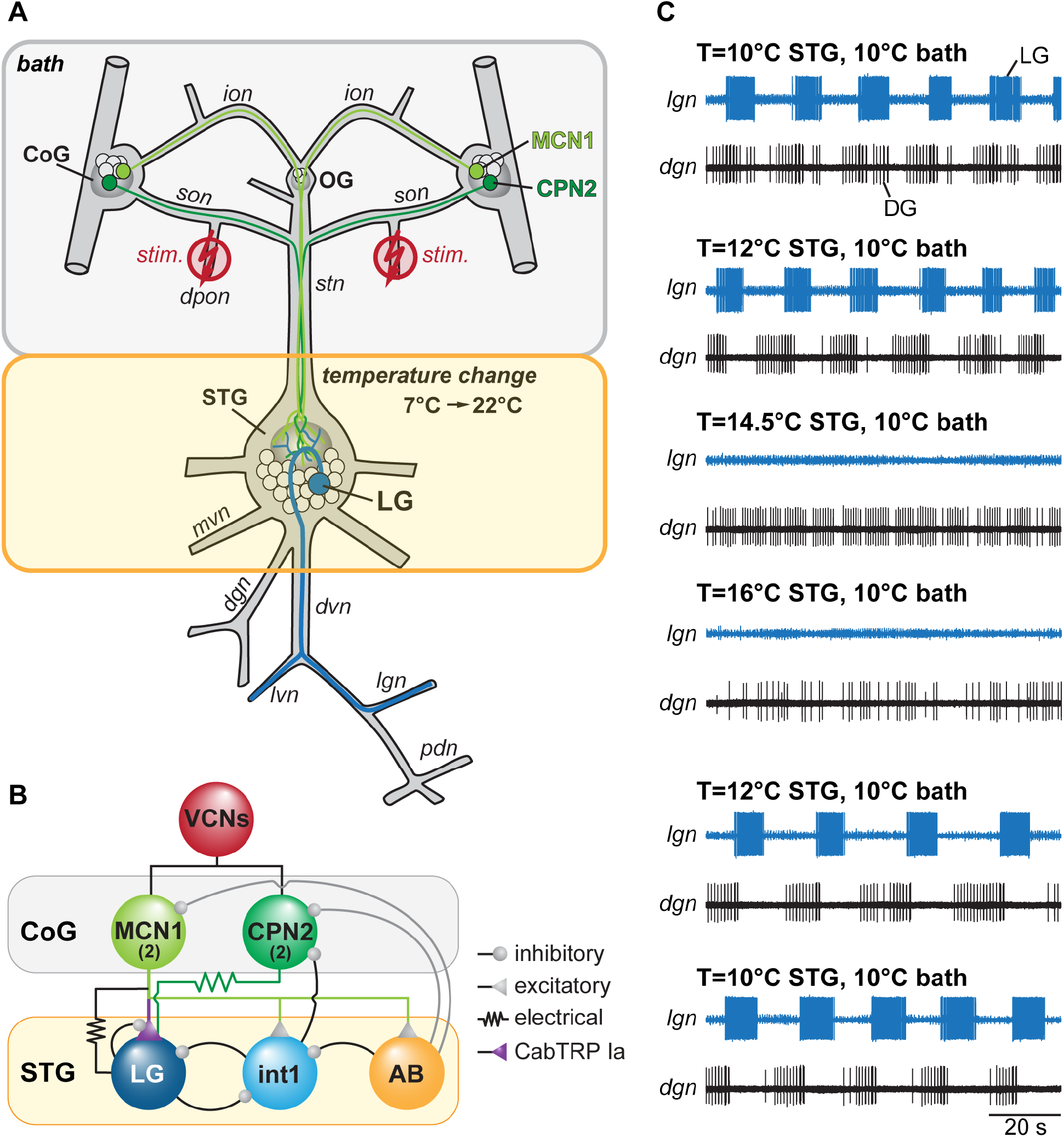
Spontaneously active gastric mill rhythms crash when only the STG is heated. **(A)** Schematic of the STNS illustrating the experimental approach. A split-bath approach was used to selectively heat either the STG (yellow), or its extrinsic inputs including the OG and the CoGs (bath, grey), which house the descending projection neurons MCN1 and CPN2. Gastric mill activity was assessed by recording the LG neuron (blue) extracellularly on the *lgn*. Except for the experiments shown in Figure 1C, gastric mill rhythms were initiated by activating the ventral cardiac neurons (VCNs) through stimulation of the paired *dpons* (‘stim.’, red). VCN activity leads to a long-lasting activity of MCN1 and CPN2 *(Beenhakker and Nusbaum, 2004)*. **(B)** Simplified wiring diagram of the VCN influence on MCN1, CPN2, the gastric mill (LG and Int1), and pyloric (AB) neurons. LG (purple) receives excitatory input from MCN1 via release of CabTRP Ia. Neuron copy numbers are in parentheses. For complete circuit diagram see *Powell et al. (2021)*. **(C)** Example of a crash of spontaneous gastric mill activity at elevated temperature. Shown are extracellular recordings of the *lgn* and *dgn* nerves at 10, 12, 14.5, and 16°C STG temperature. Bath temperature was kept constant at 10°C. The rhythm failed when STG temperature was elevated to 14.5°C or 16°C, but could be recovered by cooling the STG back to 12°C, and 10°C. ***Legend:*** CoG: commissural ganglion; OG, esophageal ganglion; STG, stomatogastric ganglion; LG, lateral gastric neuron; DG, dorsal gastric neuron; PD, pyloric dilator neuron; MCN1, modulatory commissural neuron 1; CPN2, commissural projection neuron 2; *dpon*, dorsal posterior esophageal nerve; *ion*, inferior esophageal nerve; *son*, superior esophageal nerve; *stn*, stomatogastric nerve; *dgn*; dorsal gastric nerve; *mvn*, medial ventricular nerve; *dvn*, dorsal ventricular nerve; *lgn*, lateral gastric nerve; *lvn*, lateral ventricular nerve; *pdn*, pyloric dilator nerve.

The temperature responses of these two rhythms are well characterized. The pyloric rhythm is remarkably robust and remains functional over a range of more than 20°C *(Haddad and Marder, 2018; Soofi et al., 2014; Tang et al., 2010)*, even when the pyloric central pattern generator is isolated from the animal. For the episodic gastric mill rhythm, extrinsic neuromodulation is critical for sustaining neuronal activity at elevated temperatures *(DeMaegd and Stein, 2021; Städele et al., 2015)*. Specifically, as the temperature rises even by a few degrees Celsius, LG leak conductance increases, shunting synaptic signal propagation and stopping spike activity. However, the substance P-related neuropeptide CabTRP Ia (*Cancer borealis* tachykinin-related peptide Ia) counterbalances the increase in LG’s leak and can sustain LG activity during temperature perturbation when bath-applied or neuronally released. The temperature range over which this mechanism functions is unclear. Neuromodulators, and neuropeptides in particular, have been implicated in supporting robust activity in several pattern-generating networks *(Ellingson et al., 2021; Gray et al., 1999; Mutolo et al., 2010; Shi et al., 2021; Städele et al., 2015; Zhao et al., 2011; Zhu et al., 2018)*. It is thus conceivable that extrinsic neuromodulation is a general mechanism to bestow temperature robustness to neural oscillators.

*Powell et al. (2021)* demonstrated that the gastric mill rhythm, when elicited by stimulation of the mechanosensory ventral cardiac neurons (VCN), is similarly temperature-robust as the pyloric rhythm. In addition, the integer coupling between the gastric and pyloric rhythms seems to prevail at elevated temperatures as well. However, several questions remain. Importantly, stimulation of the VCNs recruits a set of descending modulatory projection neurons whose activity is necessary to activate the gastric mill rhythm. The experiments by *Powell et al. (2021)* could not determine whether this descending modulation enables the temperature robustness of the gastric mill CPG or if the gastric mill CPG, like the pyloric CPG, is bestowed with intrinsic temperature robustness.

Furthermore, whether modulatory projection neurons extrinsic to the STG enable or influence the gastric mill integer coupling across temperature is unknown. In our study, we separated temperature effects on extrinsic projection neurons from those on the gastric mill and pyloric CPGs in the STG. We achieved this by selectively and differentially changing the temperature of the STG and the projection neurons. This enabled us to provide a mechanism for the temperature robustness of the VCN gastric mill rhythm and to determine the impact of extrinsic neuropeptide modulation on the integer coupling observed by *Powell et al. (2021)*.

## Results and Discussion

### Spontaneous gastric mill rhythms are not temperature robust without neuromodulatory compensation

The episodic gastric mill rhythm requires activation through descending projection neurons. These projection neurons are extrinsic to the stomatogastric ganglion (STG) that houses the gastric mill and pyloric CPGs. They are activated by sensory stimuli that signal food availability *(Beenhakker et al., 2004; Hedrich et al., 2009). Powell et al. (2021)* recently demonstrated that the gastric mill rhythm elicited by the mechanosensory VCN shows considerable robustness against temperature changes when both the gastric mill CPG and its extrinsic input neurons in upstream ganglia were heated. Our previous experiments show that another version of the gastric mill rhythm - elicited by stimulation of the modulatory commissural neuron 1 (MCN1) - is profoundly temperature-sensitive when neuromodulatory input from upstream projection neurons is suppressed and only the gastric mill CPG in the STG is heated *(Städele et al., 2015)*. Whether such temperature robustness for other versions of the gastric mill rhythm or just the MCN1 rhythm remains unclear.

To address this, we independently modified the temperature of the STG circuits and the projection neurons in the upstream esophageal (OG) and commissural ganglia (CoGs; Fig. 1A). For starters, we measured temperature responses of spontaneously active gastric mill rhythms. Figure 1C shows the result of such an experiment where both the STG and the upstream ganglia were kept at 10°C (top traces). When we selectively increased the temperature of the STG gastric mill CPG network but kept upstream ganglia at 10°C, we found that the rhythm continued at 12°C but ceased at 14.5°C. Increasing the STG temperature further to 16°C did not reestablish the rhythm. However, when we lowered the STG temperature back to 12°C, the rhythm reappeared and continued at 10°C. We observed similar behaviors in all preparations tested (N=5), with spontaneous gastric mill rhythms crashing at temperatures below 19°C. Together, our findings suggest that the broad temperature robustness of spontaneous gastric mill rhythms depends at least partly on heating the upstream ganglia and thus on factors extrinsic to the CPG network in the STG. This conclusion is consistent with our previous results showing that extrinsic peptide modulation bestows temperature robustness of the MCN1 gastric mill rhythm *(Städele et al., 2015)*. Given that spontaneous gastric mill rhythms almost never are generated by MCN1 alone, this indicates that extrinsic modulation is also a critical factor in sustaining other types of gastric mill rhythms.

### VCN gastric mill rhythms are not temperature robust without neuromodulatory compensation

To further investigate the impact of extrinsic modulatory factors, we stimulated the VCN via the *dpon* nerve (Figs. 1A, B). VCN stimulation activates the projection neurons MCN1 and CPN2 (commissural projection neuron 2), which in turn drive the gastric mill rhythm *(Beenhakker and Nusbaum, 2004; Blitz et al., 2004)*. We used identical stimulus protocols as *Powell and colleagues (2021)*. Consistent with previous data, stimulations elicited long-lasting (>15 minutes) gastric mill rhythms at control temperature (9 to 10°C). We then selectively altered STG temperatures (7 to 22°C, in 3°C increments), stimulated again, and tested whether gastric mill rhythms could be elicited. If a gastric mill rhythm occurred, we measured characteristics as cycle period, rhythm duration, burst durations of the neurons, and their phase relationship. Unless otherwise stated, we kept the temperature of upstream extrinsic projection neurons in the CoGs constant between 9 and 10°C (control temperature) to separate temperature effects on the gastric mill CPG from those extrinsic to it. Thus, the projection neurons activated by *dpon* stimulation and driving the VCN gastric mill rhythm (MCN1 and CPN2) remained at control temperature, while the gastric mill CPG network in the STG was exposed to changing temperatures. *Beenhakker et al. (2004)* found that VCN stimulation elicits gastric mill rhythms that last for an average of 14 minutes, with substantial variability. We took a conservative approach and tested for the presence of gastric mill rhythms five minutes after the end of the stimulation.

Figure 2A shows original recordings at STG temperatures of 7 to 19°C in 3°C steps. At 13°C, the gastric mill rhythm became irregular and was essentially absent at higher temperatures. Specifically, the core CPG neuron LG showed regular rhythmicity and burst activity at 7 and 10°C. In contrast, at 13°C, there were long pauses between LG bursts, and the cycle period started to vary. At 16°C, only a single LG burst was present, and no LG bursts occurred at 19°C (=crash). The gastric mill follower neuron DG showed regular rhythmic activity at 7 and 10°C, became arrhythmic with vastly varying burst durations at 13 and 16°C, and recovered a slow rhythmic activity with varying burst durations at 19°C.

**Figure 2.**
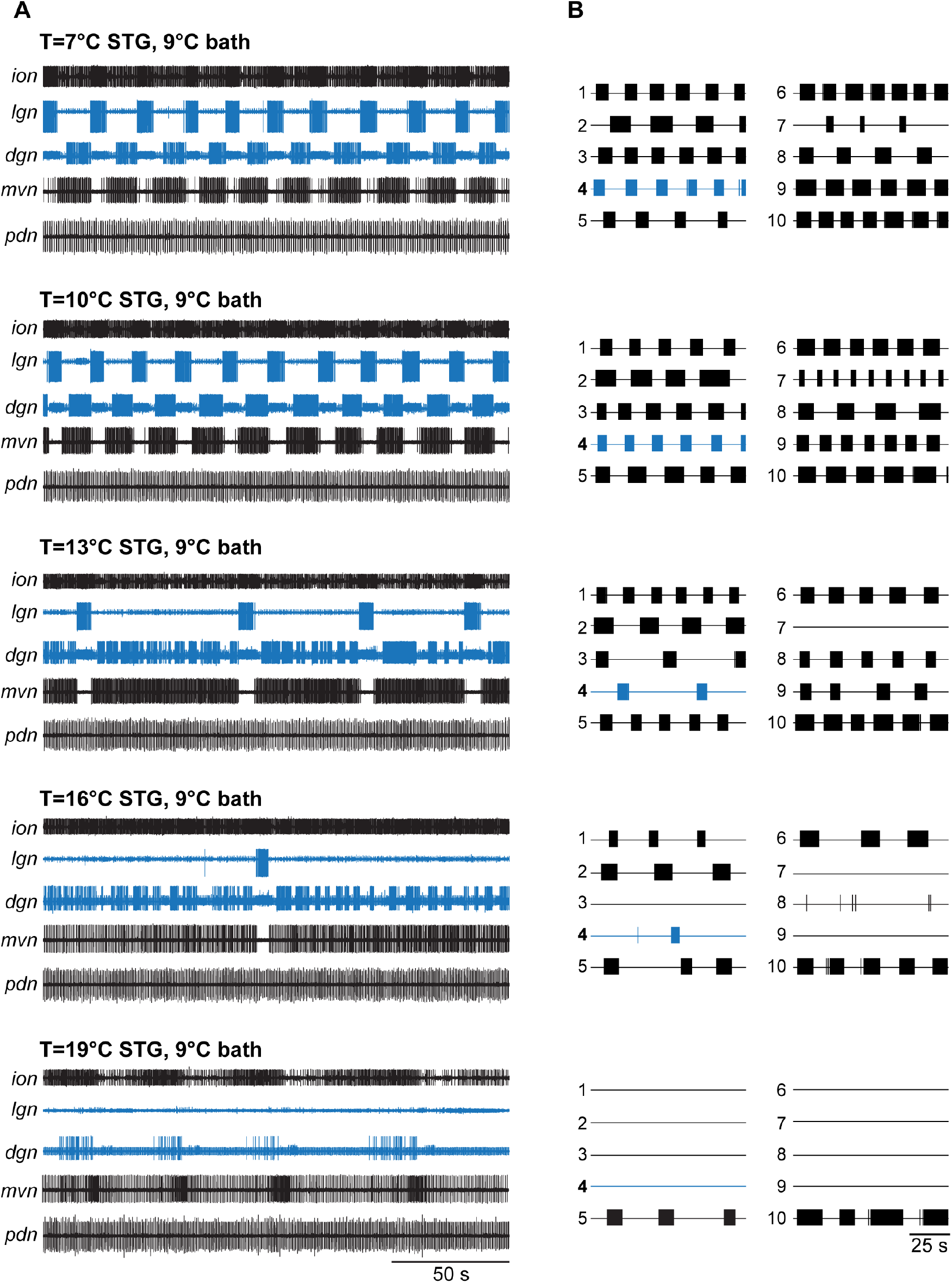
VCN gastric mill rhythms crash when only the STG is heated. **(A)** Example extracellular nerve recordings of both the gastric mill (*lgn, dgn*) and pyloric (*pdn*) rhythms at different STG temperatures. Recordings of the gastric mill neurons LG (on the *lgn* nerve) and DG (on *dgn*) are highlighted in blue. The *pdn* recoding shows the activity of the PD neurons as a representation of the pyloric rhythm. For completeness, the activity of IC and VD (both gastro-pyloric neurons) on the *mvn* was plotted as well. Note, the gastric mill (GM) neurons, which participate in the VCN rhythm, are absent from the *mvn* because the recording was taken peripherally, after all GM axons had left this nerve. For each temperature, a gastric mill rhythm was elicited with *dpon* stimulation. Bath temperature was kept constant at 9°C while STG temperature was increased. At elevated STG temperatures (16°C and 19°C), *dpon* stimulation failed to elicit a gastric mill rhythm. **(B)** Summary of all experiments (N=10 preparations). Each numbered trace represents the spike activity of LG over 100 seconds for one experiment, recorded 300 seconds after the end of the *dpon* stimulation. The blue trace corresponds to the example shown in (A).

In contrast, the pyloric rhythm continued to produce regular activity across temperatures, consistent with previously published data *(Soofi et al., 2014; Tang et al., 2010)*. Figure 2A shows this with a recording of the PD pyloric pacemaker neurons on the *pdn*.

Our data were consistent across preparations tested (Fig. 2B), although the lowest temperature at which stimulation failed to elicit a lasting rhythm varied (13°C: N=1/out of 10 preparations crashed, 16°C: N=3/10, 19°C: N=9/10, 22°C: N=10/10). In all temperature conditions, VCN stimulation elicited the typical responses during stimulation *(Beenhakker et al., 2004)*. In some cases, when the rhythms crashed, a few cycles of gastric mill activity were observed immediately after the end of the stimulation. However, this activity was short-lived and never continued. Figure 3A (red) shows the percentage of rhythms present 5 minutes after *dpon* stimulation for the various temperatures tested. For analysis, we scored rhythms as present when at least 3 LG bursts occurred within 200 seconds, no matter how irregular their occurrence was.

**Figure 3.**
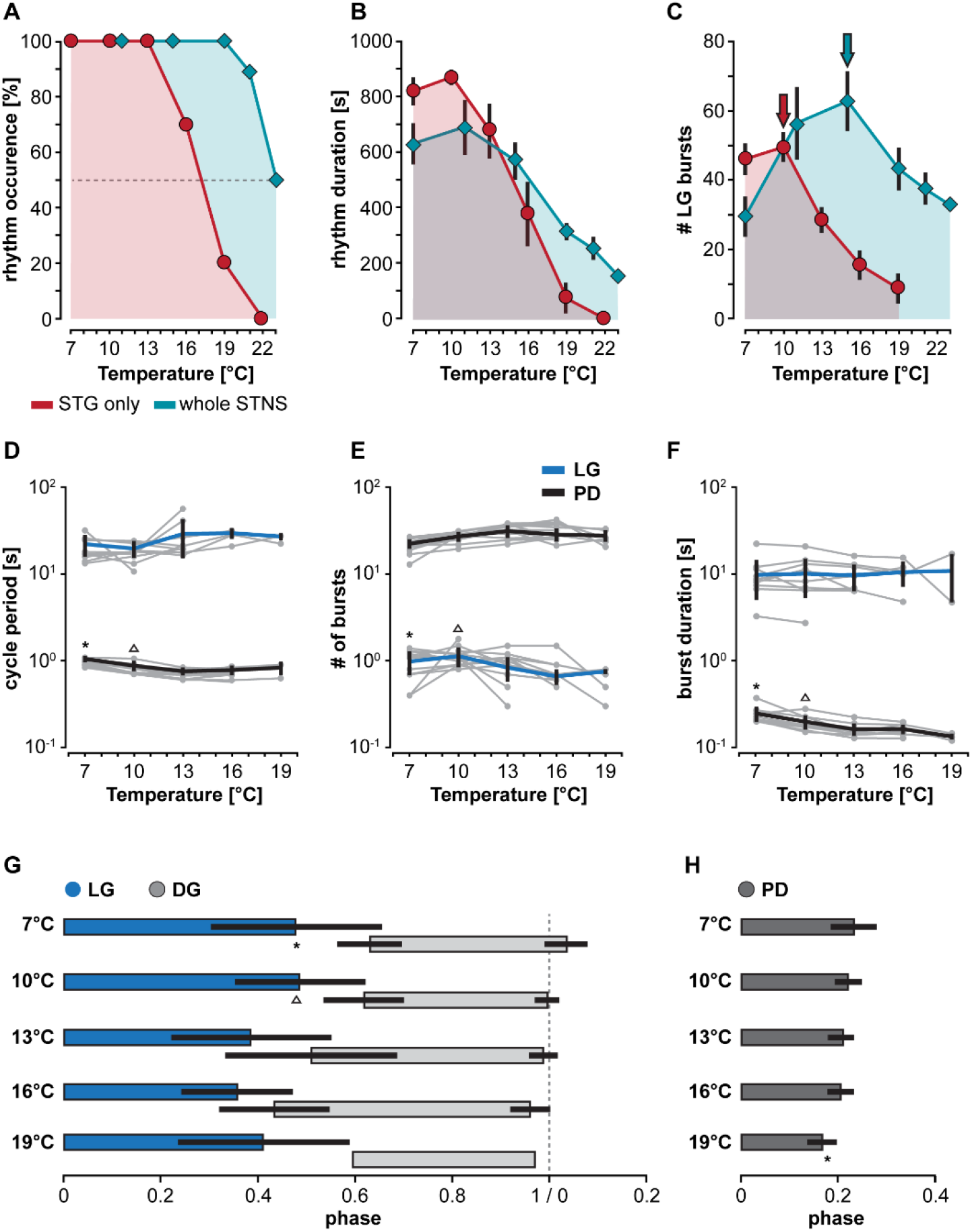
Heating extrinsic input fibers improves temperature resistance of CPGs in the STG. **(A-C)** Comparison of gastric mill rhythm parameters for two different heating condition. Red traces: STG temperature changed while CoGs were kept at 9°C. Teal traces: STG and CoG temperatures were changed together (reanalyzed data from *Powell et al. (2021)*). Dots are representing mean values ± SEM of 10 preparations. For better comparison, areas underneath the curves were colored. **(A)** Number of gastric mill rhythm occurrences at different temperatures. **(B)** Gastric mill rhythm duration, and **(C)** LG spike number at different temperatures. **(D-F)** Responses of pyloric and gastric mill rhythms when only the STG is heated. Cycle periods of LG (blue) and PD (black) are shown across temperature. Thick lines represent the mean ± SEM for 10 preparations tested. Individual experiments are depicted in gray. Dots represents mean values of data collected over 200 seconds, recorded 300 seconds after the end of the *dpon* stimulation. The gastric mill rhythm crashed at varying temperatures, visible by the decline in the number of gray dots for LG at higher temperatures. **(D)** LG and PD cycle period across temperature. Statistics: LG cycle period: not significant different, One Way RM ANOVA, N=10 (7-10°C), N=9 (13°C), N=5 (16°C), N=2 (19°C), F(4,22)=2.436, p=0.078. PD cycle period: One Way RM ANOVA, N=10 (7-16°C), N=3 (19°C), F(4,29)=42.651, p<0.001, significantly different for 7°C vs. all other temperatures (indicated with *), and 10°C vs. all others (indicated with Δ), p<0.05, Student-Newman-Keuls post-hoc test. **(E)** Number of LG and PD bursts across temperature. LG bursts: not significant different, One Way RM ANOVA, N=10 (7-10°C), N=9 (13°C), N=5 (16°C), N=2 (19°C), F(4,22)=2.582, p=0.065. PD bursts: One Way RM ANOVA, N=10 (7-16°C), N=3 (19°C), F(4,29)=11.851, p<0.001, significantly different for 7°C vs. 13° and 16°C (*). 10°C was also significantly different from 13°C (Δ), p<0.05, Student-Newman-Keuls post-hoc test. **(F)** LG and PD burst duration across temperature. LG burst duration: no significant change, One Way RM ANOVA, N=10 (7-10°C), N=9 (13°C), N=5 (16°C), N=2 (19°C), F(4,22)=1.067, p=0.396. PD burst duration: One Way RM ANOVA, N=10 (7-16°C), N=3 (19°C), F(4,29)=22.752, p<0.001, significant differences for 7°C (*) and 10°C (Δ) vs. all higher temperatures, p<0.05, Student-Newman-Keuls post-hoc test. **(G)** Phase relationship and duty cycle of LG (blue), DG (light grey), and PD **(H)** across various STG temperatures. LG duty cycle (= end of its activity phase): One Way RM ANOVA, N=10 (7-10°C), N=9 (13°C), N=5 (16°C), N=2 (19°C), F(4,22)=10.242, p<0.001, significant differences for 7°C (*) and 10°C (Δ) vs. all higher temperatures, p<0.05, Student-Newman-Keuls post-hoc test. At 19°C, DG was only active in one animal. DG phase onset: significant effect of temperature (p=0.018), but no significant difference between any two conditions. One way ANOVA, N=8 (7°C), N=10 (10°C), N=9 (13°C), N=5 (16°C), N=1 (19°C), F(3,28)= 3.938, p=0.018, p>0.05 for all paired comparisons, Student-Newman-Keuls post-hoc test. DG phase end: One way ANOVA, N=8 (7°C), N=10 (10°C), N=9 (13°C), N=5 (16°C), N=1 (19°C), F(3,28)=5.395, p=0.005, significant differences between 7 and 16°C (*), p<0.05, Student-Newman-Keuls post-hoc test. DG duty cycle: no significant effect of temperature, One way ANOVA, N=8 (7°C), N=10 (10°C), N=9 (13°C), N=5 (16°C), N=1 (19°C), F(3,28)= 2.786, p=0.059. PD: One Way RM ANOVA, N=10 (7-16°C), N=3 (19°C), F(4,29)=6.019, p=0.001, significant differences for 19°C vs. all others (*), p<0.05, Student-Newman-Keuls post-hoc test.

The crash of the gastric mill rhythm at relatively low temperatures contrasts with findings from *Powell et al. (2021)*, where the CoGs were heated to the same temperature as the STG. For comparison, we thus reanalyzed publicly available data from *Powell et al. (2021)* and plotted them next to ours (Fig. 3A, teal). The difference is quite striking in that all rhythms where the bath (and thus also the CoGs) was kept at control temperatures crashed at lower or equal temperatures than the rhythms where STG and CoG temperature were the same. This difference was also reflected in rhythms generally ending earlier after the *dpon* stimulation when only the STG was heated (Fig. 3B), with correspondingly fewer total LG bursts (Fig. 3C). We noted that independent of the heating condition (entire preparation vs. STG only), the total number of LG bursts generated after *dpon* stimulation reached a peak before decreasing at higher temperature. When only the STG was heated, the peak occurred at 10°C in comparison to 16°C when the entire preparation was heated (Fig. 3C, arrows).

### Extrinsic factors contribute to the temperature responses of pyloric and gastric mill rhythms

To further characterize the temperature response of the gastric mill rhythm, we analyzed the gastric mill cycle period, the number of bursts per time, LG burst duration, and LG duty cycle (over 200 seconds). We noted a pronounced difference to data from *Powell et al. (2021)*. In our experiments, where only the STG was heated, the gastric mill period did not decrease dramatically with temperature (Fig, 3D). Instead, the cycle period tended to increase with temperatures over 10°C, albeit not significantly. Similarly, the number of LG bursts per time remained unchanged (Fig. 3E). LG burst duration was also constant despite the increasing temperature (Fig. 3F). We found, however, that because of the trend towards increasing cycle periods and constant burst durations, LG’s duty cycle in the rhythm decreased at 13°C and above (Fig. 3G). The decrease in LG duty cycle was accompanied by an increase in the duty cycle of the dorsal gastric (DG) neuron (Fig. 3G), LG’s functional antagonist. With respect to the behavioral output of the rhythm, this shortens the protraction phase and extends the retraction phase. In intact animals, even small changes in LG burst activity are associated with changes in the movement of the stomach teeth *(Diehl et al., 2013; Heinzel, 1988a, 1988b)*, suggesting that changes in the LG duty cycle at higher temperatures will lead to behavioral changes. Together, our data demonstrate that dissociating temperature effects on the CPG from those of upstream ganglia substantially lowered the robustness of the gastric mill rhythm against temperature perturbation, suggesting that factors extrinsic to the CPG provide a functionally relevant component to temperature compensation.

We further noted that the variability of the rhythm did not increase significantly when approaching the crash temperature. The original recordings in Figure 2 already show this trend in that there was either regular rhythmic burst activity or no burst activity at all. We also did not find quantitative differences in rhythm variability when we compared the co-efficient of variation of the gastric mill cycle period at 10°C and at the highest temperature at which a rhythm could still be elicited (10°C: 0.11±0.07, pre-crash temperature: 0.17±0.13; paired t-test, N=10, P = 0.134).

While the pyloric rhythm continued to function in all of our experiments, even at temperatures where the gastric mill rhythm had crashed, we noted more differences to previous experiments where CoGs and STG were heated together *(Powell et al., 2021)*. Most obviously, although there was a significant decrease in the pyloric cycle period at temperatures of 13°C and above (Fig. 3D), this change seemed modest when compared to *Powell et al. (2021)*. To determine the pyloric cycle period, we used the bursts of the pyloric dilator neurons on the *pdn* and measured the time between their onsets. In agreement with the small effect on the cycle period, there was no consistent change in the total number of PD bursts (Fig. 3E). This suggests that some aspects of the speeding up of the pyloric rhythm that is observed in intact animals *(Soofi et al., 2014)* and during heating of the entire nervous system *(Haddad and Marder, 2018; Powell et al., 2021; Tang et al., 2012, 2010)* are likely to be mediated by extrinsic factors.

We also found similarities to previously published data. For example, like previously described for varying temperatures both *in vitro* and *in vivo (Soofi et al., 2014; Tang et al., 2010)*, the duty cycle of the pyloric pacemaker neurons remained mainly constant throughout the temperature range tested. Accordingly, PD burst duration (Fig. 3F) mirrored the pyloric cycle period in that it shortened significantly at temperatures of 13°C and above, which together lead to a primarily constant duty cycle. Only when the temperature reached 19°C, there was a significant shortening of the duty cycle (Fig. 3H). Taken together, our data demonstrate that factors extrinsic to the STG, and thus to the pyloric and gastric mill CPGs, contribute to the temperature responses of both rhythms. For the gastric mill rhythm, neurons residing outside the STG appear to provide a significant component to the substantial temperature robustness of the rhythm.

### VCN gastric mill rescue is accompanied by temperature-induced increase in modulatory projection neuron activity

In previous studies *(DeMaegd and Stein, 2021; Städele et al., 2015)*, we have shown that extrinsic modulation enables the temperature robustness of a specific version of the gastric mill rhythm, elicited by the paired descending modulatory projection neuron MCN1. Specifically, the peptide co-transmitter of MCN1, CabTRP Ia, reduces shunting in LG to counterbalance an excessive increase in LG leak currents. CabTRP Ia is released increasingly with higher MCN1 firing frequencies *(Stein et al., 2007)*. The increased MCN1 firing frequency strengthens the robustness of the gastric mill rhythm against temperature perturbations *(Städele et al., 2015)*. MCN1 and CPN2 drive the VCN version of the gastric mill rhythm investigated here. MCN1 releases only one peptide that affects LG (CabTRP Ia) *(Nusbaum and Beenhakker, 2002; Stein et al., 2007)*. While CPN2’s neurotransmitters are unknown, most of its actions in the STG are mediated through electrical synapses. CPN2’s effects on LG are weak compared to MCN1 *(Stein et al., 2007)* but lead to the recruitment of additional gastric mill neurons into the rhythm *(Beenhakker and Nusbaum, 2004)*. While multiple projection neurons are active during a VCN rhythm, MCN1 is the main contributor to LG excitation. This suggests that temperature and compensation effects may be similar to the MCN1 gastric mill rhythm. We thus hypothesized that the temperature robustness of the VCN gastric mill rhythm is enabled, at least in part, by extrinsic peptide modulation from MCN1.

We thus tested whether temperature-induced up-regulation in projection neuron firing frequency is sufficient to prevent the termination of the VCN gastric mill rhythm at elevated temperatures. Like in our previous experiments, we stimulated the *dpon* to elicit a VCN gastric mill rhythm while we heated the STG to different temperatures. We then determined the crash temperature, i.e., the temperature at which stimulation failed to elicit a gastric mill rhythm. The ganglia extrinsic to the STG were then heated to the crash temperature. The *dpon* was stimulated again with both, the STG and the CoGs, at crash temperature. Figure 4A shows a representative recording of LG at 16°C (4A, i), where the rhythm could still be elicited, and then at a crash temperature of 19°C (ii), where the rhythm failed when only the STG was heated. After additionally heating the CoGs to 19°C (iii), however, the gastric mill rhythm could be recovered with *dpon* stimulation. In all preparations tested (N=5, Fig. 4B), heating of the STG and CoGs together led to a rescue of gastric mill activity at crash temperature.

**Figure 4.**
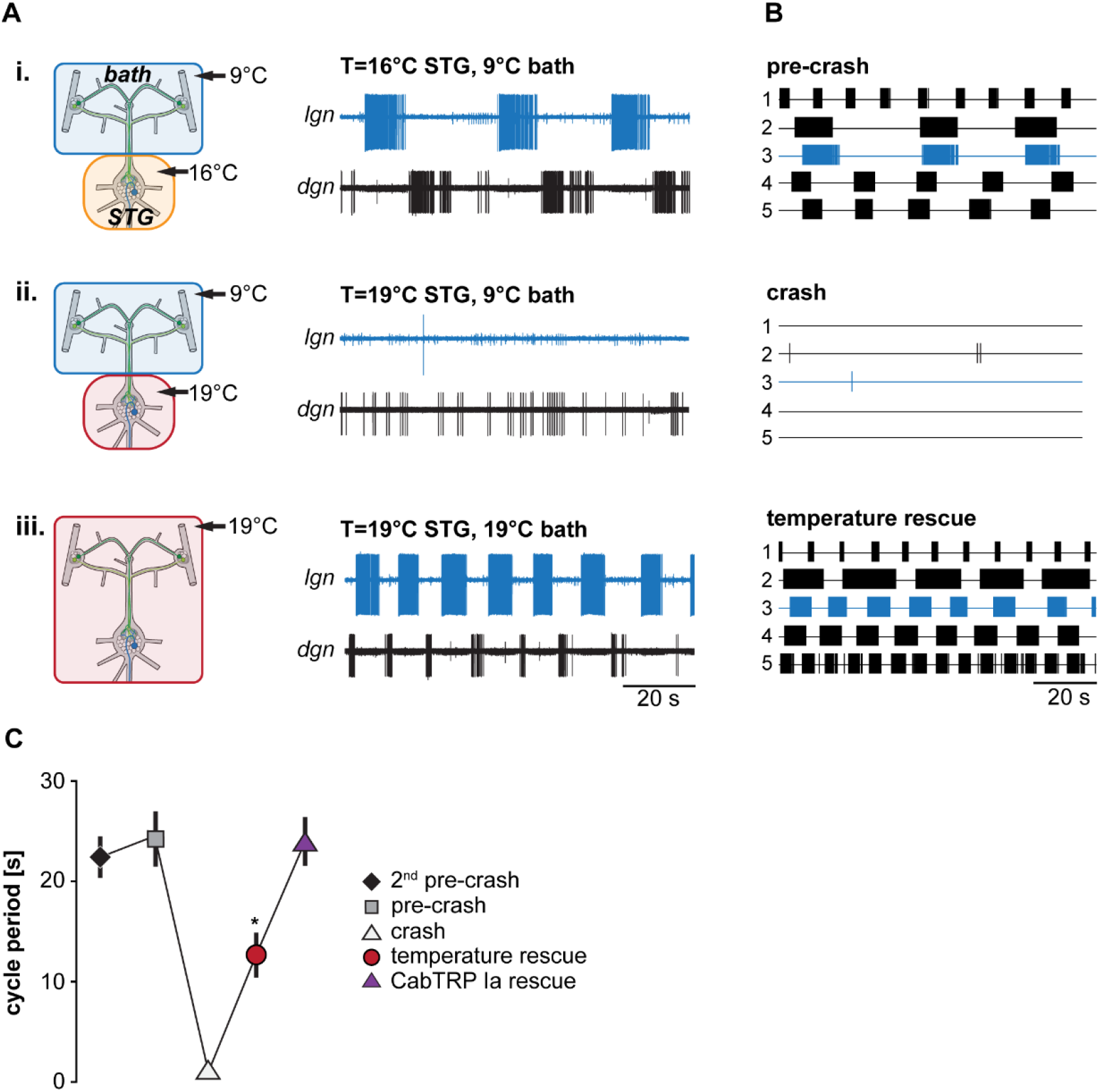
Matching STG and CoG temperature prevents gastric mill rhythm crash. **(A)** Schematic representation of the temperature conditions (left) and the corresponding activity of LG and DG for one preparation (right). **(B)** Summary of all experiments (N=5). Each trace represents the spike activity of LG over 100 seconds for an individual experiment, recorded 300 seconds after the end of the *dpon* stimulation. Blue traces correspond to data shown in A. Crash temperatures varied between experiments. Conditions are labeled as pre-crash, crash, and temperature rescue. **(C)** Comparison of LG cycle period for the specified conditions. Shown are mean values ± SEM of all preparations (N=10). For completeness, LG cycle period during bath application of CabTRP Ia (Fig. 6) was added to this plot (N=9). LG cycle period was significantly decreased during temperature rescue compared to the 2^nd^ pre-crash, pre-crash, and CabTRP Ia rescue (indicated with *). One Way ANOVA, F(3,30)=55.972, p=0.046, p<0.05 for paired comparisons, Student-Newman-Keuls post-hoc test.

We further noted that the rhythm sped up during this rescue, with a significantly shorter cycle period than at the last temperature before the crash (pre-crash, Fig.4C). Our results demonstrate that heating the CoGs rescued the rhythm and enabled faster cycle periods. This result is consistent with the observation of *Powell et al. (2021)* that increasing temperatures caused faster rhythms when STG and CoGs were heated together. In summary, our data suggest that a heat-activated factor extrinsic to the STG is necessary to enable temperature robustness of the gastric mill rhythm.

Our previous work had indicated that the activity of the CoG projection neurons MCN1 increases with temperature *(Städele et al., 2015)*, albeit in different experimental conditions. We hypothesized that MCN1 and its co-transmitters contribute to the temperature robustness of the VCN gastric mill rhythm and predicted that its activity increases when both the STG and CoGs are warmed to the same temperature. Conversely, we also predicted that MCN1 activity remains unchanged when only the STG is heated, ultimately leading to a crash of the gastric mill rhythm. To test these predictions, we investigated MCN1’s firing frequency after *dpon* stimulation in the different STG temperature conditions (STG heated vs. entire preparation heated). Figure 5A shows original recordings of the two MCN1 neurons on the left and right inferior esophageal nerves (*ion*) and the corresponding MCN1 firing frequencies for three different temperature conditions. LG activity was monitored extracellularly on the *lgn*. At pre-crash temperature (STG 16°C, CoGs 9°C, Fig. 5A, i), VCN stimulation elicited a regular, slow gastric mill rhythm. MCN1’s firing frequency varied rhythmically, with the highest frequencies coinciding with the LG burst onset and lowest frequencies immediately following the LG burst end. This progression of firing frequencies is reminiscent of previously published data *(Beenhakker and Nusbaum, 2004)* and partly depends on feedback from the gastric mill CPG to the commissural projection neurons *(Blitz, 2017)*. Figure 5A, ii shows MCN1 firing frequencies after the rhythm had crashed (STG 19°C, CoGs 9°C).

**Figure 5.**
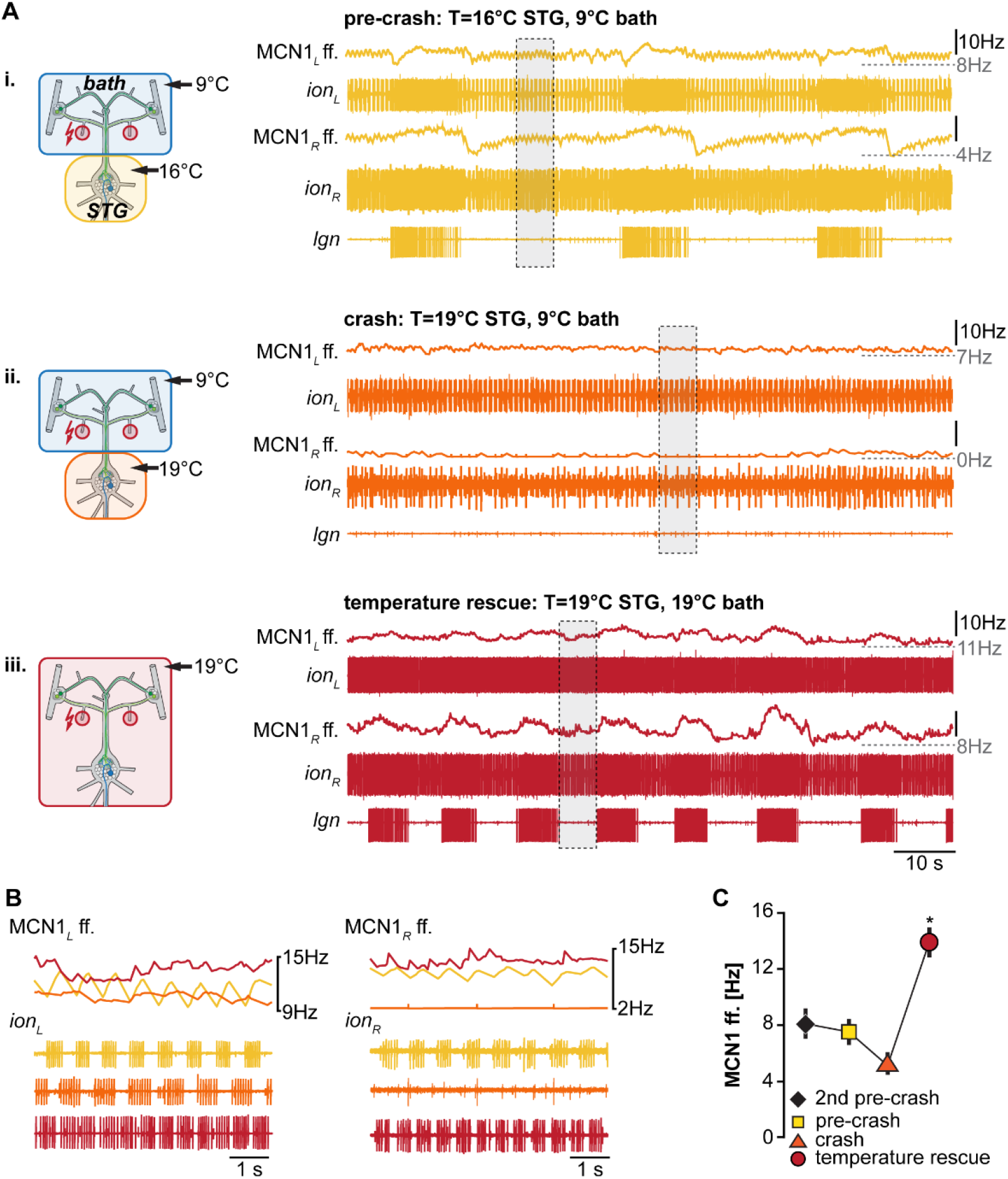
Matching STG and CoG temperature increases MCN1 firing frequency. **(A)** Schematic representation of the temperature conditions (left) and the corresponding activity of MCN1 for one preparation (right). Firing frequencies for each of the two MCN1 neurons are shown. L corresponds to the left CoG and *ion*, and R to the right. Traces above each *ion* recording show the mean MCN1 firing rate (sliding window of 1 second). The bottom recording shows the *lgn*, and the LG neuron and gastric mill rhythm. **(B)**. Top: Comparison of MCN1 mean firing frequency for both MCN1 neurons (left and right). Bottom: expanded view of the *ion* recordings depicted in A with gray boxes. Colors represent the temperature conditions shown in A. **(C)** Comparison of mean MCN1 firing frequency for all experiments (N=4 for temperature rescue, N=8 all other conditions). MCN1_L_ and MCN1_R_ frequencies were averaged. The MCN1 firing frequency in the temperature rescue condition was significantly increased in comparison to all other conditions (indicated with *). One Way ANOVA, F(3,24)=5.102, p<0.001, p<0.05 for all paired comparisons, Student-Newman-Keuls post-hoc test.

Note that MCN1 firing frequencies were similar to the lowest frequencies of the pre-crash temperature. However, LG bursts were absent, indicating that MCN1 could not recruit LG, so the gastric mill rhythm crashed. In fact, LG was silent, stopping CPG oscillation. As a consequence of the missing CPG feedback to the commissural projection neurons, rhythmic increases in MCN1 firing frequency were also absent in this condition, further lowering the chances of recruiting LG. Bringing the CoGs to the same temperature as the STG circuits (both at 19°C) caused an increase in MCN1 firing frequencies and restored the rhythm (Figure 5A, iii). Figure 5B compares MCN1 firing frequencies for the areas highlighted with gray boxes in Figure 5A, separated for the left and right MCN1. Examples were taken from the LG interburst period, i.e., when MCN1 activities were lowest during rhythms. MCN1 activity was lowest at the crash temperature, and it was highest in the temperature rescue condition. We found this to be true for all preparations tested (N=4). Figure 5C shows the averaged summary of all experiments, including two temperatures before the rhythm crashed. There was no apparent effect on MCN1 firing frequency at the two temperatures before the crash. The lowest MCN1 activity was consistently observed at crash temperature, in accordance with the lack of gastric mill rhythm and STG circuit feedback to the CoGs. Importantly, when the CoGs were heated to the crash temperature (temperature rescue), MCN1 activity increased significantly, and the rhythm was restored. These data are consistent with the idea that increased MCN1 activity rescued the gastric mill rhythm.

### Neuromodulator application rescues VCN gastric mill rhythms

MCN1’s neuropeptide co-transmitter CabTRP Ia has previously been indicated to provide temperature-robustness to LG’s activity *(Städele et al., 2015)*. These experiments, however, had been done for the MCN1-only gastric mill rhythm. We suspected that CabTRP Ia might have a similar effect on the VCN gastric mill rhythm. To test this, we again elicited gastric mill rhythms by stimulating the *dpons* (Fig. 6A, i), increased the STG temperature until no rhythm could be elicited anymore (Fig. 6A, ii), and then bath applied CabTRP Ia (1μM) selectively to the STG. The CoGs were always kept at control temperature (9°C). Applying CabTRP Ia by itself did not alter LG activity, i.e., no gastric mill rhythm was elicited by simply adding CabTRP Ia (Fig. 6A, iii). However, stimulating the *dpon* in CabTRP Ia restored the gastric mill rhythm at crash temperature (Fig. 6A, iv), resembling the situation in the MCN1-only rhythm *(Städele et al., 2015)* and the temperature rescue of the VCN rhythm (Fig. 5A, iii). In all preparations tested (N=7), CabTRP Ia application was sufficient to restore the VNC rhythm at elevated temperatures (Fig. 6B). We noted however that unlike in the temperature rescue experiments in which the CoGs and STG were heated together (Fig. 5A, B), rhythms in CabTRP Ia showed no significant change in cycle period to the pre-crash rhythms in saline (Figure 4D). This suggested that the continuous presence of MCN1’s co-transmitter is insufficient to establish the temperature-induced increase in gastric mill rhythm speed when the whole nervous system is heated.

**Figure 6.**
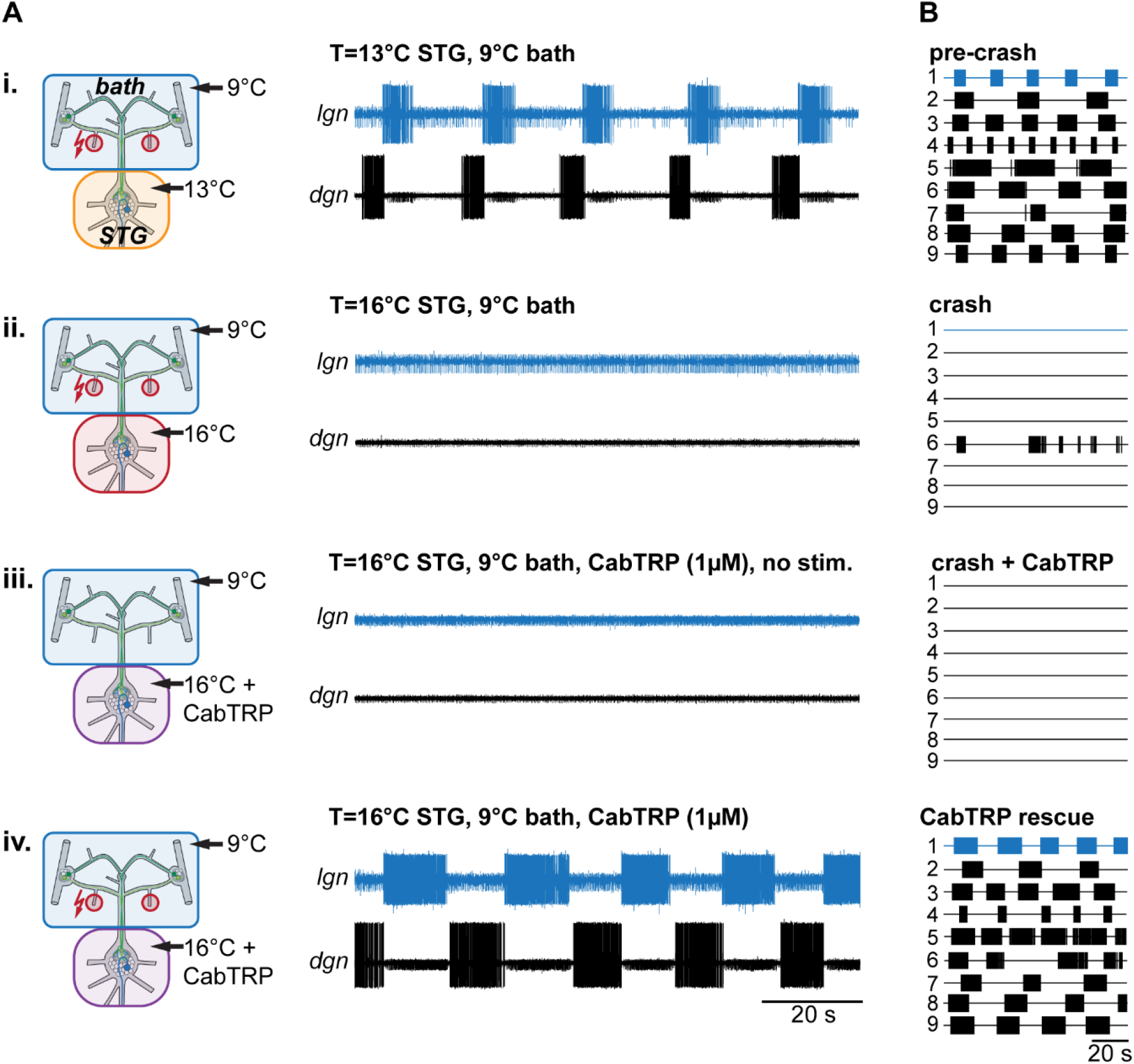
Rhythms can be recovered by bath application of MCN1’s peptide co-transmitter CabTRP Ia. **(A)** Schematic representation of the temperature conditions (left) and the corresponding activity of LG and DG for one preparation (right). **(B)** Summary of all experiments (N=9). Each numbered trace represents the spike activity of LG over 100 seconds for one experiment, recorded 300 seconds after the end of the *dpon* stimulation. The blue traces correspond to the experiment shown in B.

In summary, our data show that MCN1’s co-transmitter CabTRP Ia can stabilize the VCN gastric mill rhythm over a broad temperature range. Our data further suggest that increased MCN1 activity and CabTRP Ia release are critical components of the temperature robustness of gastric mill rhythms that involve MCN1. The temperature compensation by CabTRP Ia is mediated through the neuromodulator-induced current I_MI_ *(DeMaegd and Stein, 2021)*. I_MI_ can be activated by several other neuropeptides and even muscarinic agonists *(Swensen and Marder, 2000)*. Crustacean cardioactive peptide (CCAP), for example, is a hormone that induces I_MI_ in LG *(DeLong et al., 2009)* and can also confer temperature robustness to the gastric mill rhythm *(DeMaegd and Stein, 2021)*. It is thus conceivable that extrinsic neuropeptide modulation sustains all gastric mill rhythms, even those independent of MCN1, and is likely to bestow considerable temperature robustness to the gastric mill rhythm *in vivo*.

Similar rescue mechanisms may also contribute to the robustness (or lack thereof) in other circuits and animals. Independently of the specific circuit, elevated temperature will, in almost all cases, lead to an increase of ionic conductance and leak currents. Membrane shunting because of increased leak currents has long been known to play essential roles in the regulation of excitability *(Brickley et al., 2007; Rekling et al., 2000)*, switching neuronal activity states (Gramoll et al., 1994), and in the control of network oscillations *(Blethyn et al., 2006; Cymbalyuk et al., 2002; Koizumi et al., 2008; Zhao et al., 2010)*. Consequently, modulation of leak currents by neurotransmitters has been proposed to contribute to the regulation of neuronal excitability *(Aller et al., 2005; Kim, 2005; Pratt and Aizenman, 2007)*. There are also several examples where neuromodulators, and neuropeptides in particular, have been implicated in supporting robust activity in several pattern-generating networks *(Ellingson et al., 2021; Gray et al., 1999; Mutolo et al., 2010; Shi et al., 2021; Zhao et al., 2011; Zhu et al., 2018)*. Increasing the robustness of a circuit through extrinsic modulation may be especially meaningful when perturbations occur rapidly, like in intertidal species that experience substantial temperature variations within a few minutes *(Stein and Harzsch, 2021)*.

### Integer coupling is determined by STG circuit architecture and does not depend on extrinsic neuromodulation

*Powell et al. (2021)* have recently shown that integer coupling between the pyloric and the VCN gastric mill rhythm is robust across temperatures (up to 25°C). LG bursts are initiated when Int1, the half-center antagonist of LG, is inhibited by the pyloric pacemaker neuron AB (Fig. 1B). AB bursting leads to a disinhibition of LG upon which the LG burst is initiated after a short delay. As a consequence of the pyloric-timed disinhibition of LG, each gastric mill cycle is comprised of an integer multiple of pyloric cycles.

LG’s pyloric-timed disinhibition from Int1 is the primary mechanism that determines the timing of the MCN1 gastric mill rhythm *(Bartos et al., 1999)*. Without this disinhibition, LG bursts would not be timed by the pyloric CPG, and integer coupling would be absent. There are also indications that integer coupling may be present *in vivo (Hedrich et al., 2011; Yarger and Stein, 2015)*. While not explicitly mentioned, work from *Powell et al. (2021)* suggests that LG disinhibition is also the primary mechanism determining the timing of the VCN rhythm, no matter the temperature. Integer coupling between the pyloric and the gastric mill rhythm is caused by the circuit architecture of the CPG’s located in the STG and is thus CPG-intrinsic.

Consequently, integer coupling is likely independent of neuromodulatory input and maintained across temperatures as long as both rhythms are active. To test this, we again specifically changed the temperature of only the STG circuits and measured the coupling strength of the two rhythms following the same procedures as *Powell et al. (2021)*. In short, we analyzed the relationship between gastric mill (LG) and pyloric (PD) cycle period (Figure 7A). Like in *Powell et al. (2021)*, most of our data tended to lie along lines with integer slope (gray lines). Only a few data points were between slopes, demonstrating that integer coupling was independent of STG temperature. Our data thus support the hypothesis that the mechanisms establishing integer coupling are intrinsic to the CPG circuits within the STG. Integer coupling does not strongly rely on extrinsic factors to be maintained when the temperature changes. However, we noted that in our data set there were fewer data points near the axes’ origins than when STG and CoGs were heated together (comparison with Fig. 5b, *Powell et al., 2021*). In our experiments where only the STG was heated, the gastric mill rhythm slowed down at higher temperatures while the pyloric rhythm sped up. In contrast, when STG and CoGs were heated together *(Powell et al., 2021)*, both rhythms sped up, resulting in more cycles with short cycle periods and thus more data near the axes’ origins.

**Figure 7.**
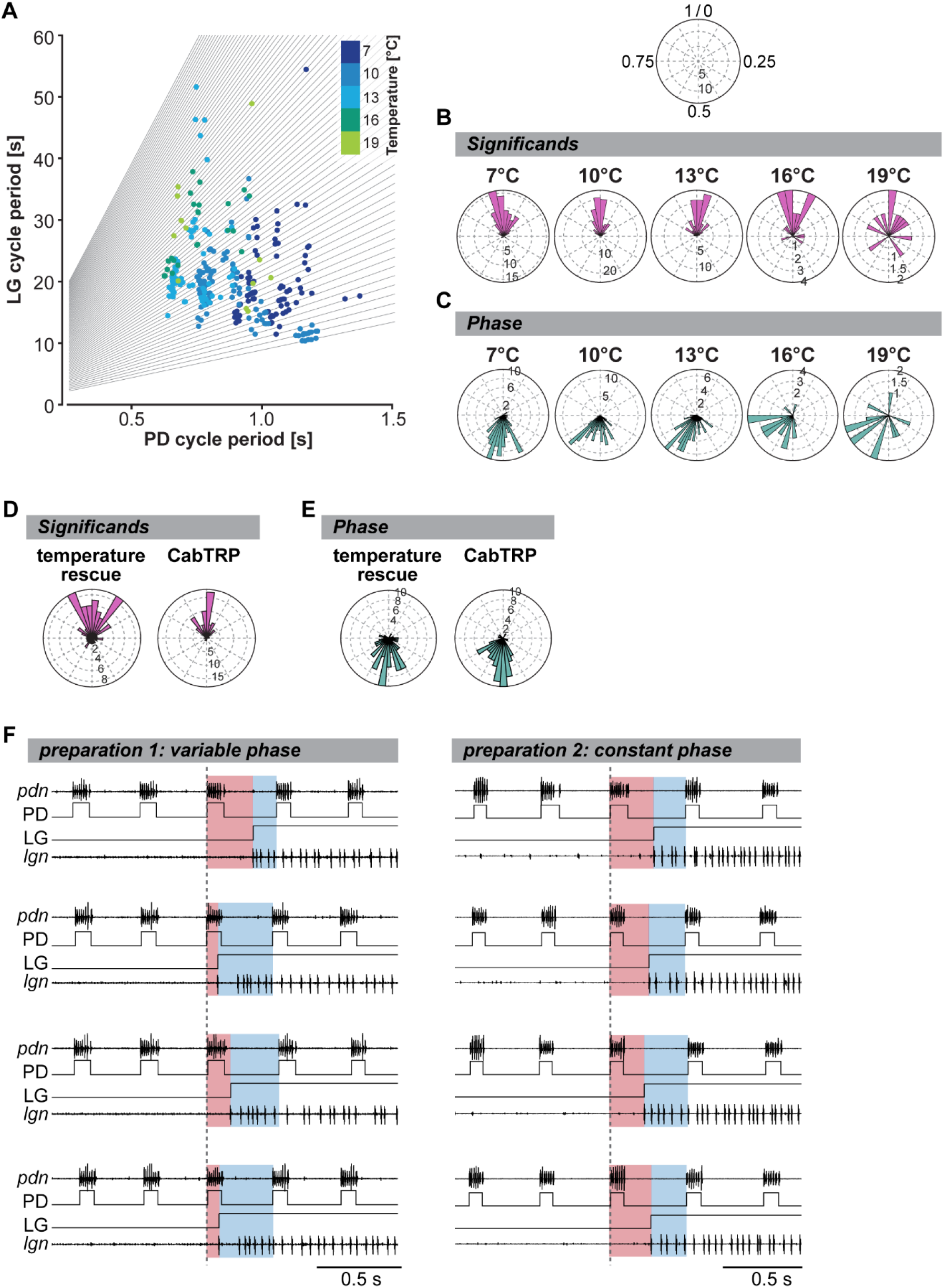
Integer coupling is always present when gastric mill and pyloric rhythms are active but weakens at elevated temperature. **(A)** LG cycle period as a function of mean PD period (= mean of all PD bursts within one LG cycle period). Gray lines represent the integer slopes and data points with integer coupling lie along these lines. N=10 preparations. **(B)** Significands for all tested preparations (N=10) shown as rose plots for various STG temperatures. Integer coupled data points will coincide at the top of the diagram (close to 1/0, see schematic at the top). Vector strengths are indicated by the radius of the plot. Analyzed were 200 seconds of recordings, 300 seconds after the last *dpon* stim. Rayleigh z Test: 7°C: Rayleigh z_65.6_, p<0.001, 10°C: Rayleigh z_83.8_, p<0.001, 13°C: Rayleigh z_49.7_, p<0.001, 16°C: Rayleigh z_12.6_, p<0.001, 19°C: Rayleigh z_3.0_, p=0.045. **(C)** Analysis of phase coupling of the LG and PD burst onset at various temperatures (N=10 preparations). Rayleigh z Test: 7°C: Rayleigh z_64.2_, p<0.001, 10°C: Rayleigh z_77.3_, p<0.001, 13°C: Rayleigh z_46.3_, p<0.001, 16°C: Rayleigh z_12.8_, p<0.001, 19°C: Rayleigh z_3.5_, p=0.027. **(D)** Significands of all gastric mill cycles and animals for the temperature rescue (N=10) and CabTRP Ia (1μM, N=9) experiments. Rayleigh z Test: temperature rescue: Rayleigh z_24.3_, p<0.001; CabTRP Ia rescue: Raleigh z_47.2_, p<0.001. **(E)** Phase coupling of the LG and PD burst onsets for the temperature rescue and CabTRP Ia (1μM) experiments. Rayleigh z Test: temperature rescue: Rayleigh z_38.5_, p<0.001; CabTRP Ia rescue: Rayleigh z_48.9_, p<0.001. **(F)** Example recordings from two preparations during temperature rescue. The LG burst is shown with a recording of the *lgn* and schematized LG bursts (LG). Four individual LG burst onsets are displayed (top to bottom) for each preparation. Pyloric cycles are shown by recordings of the PD neurons (*pdn*) and schematized PD bursts (PD). The dashed vertical line indicates the beginning of the last PD burst before the LG burst onset. The time delay between PD onset and LG onset is highlighted in red. The remainder of the pyloric cycle is marked in blue. A change in the ratio of red and blue illustrates a shift in the phase relationship between PD and LG onset. The phase relationship varied for the preparation shown on the left but not for the one on the right. **Figure supplement S1:** Data from Powell et al. (2021), reanalyzed.

To further quantify integer coupling, we calculated the significand according to *Powell et al. (2021)*. The significand provides a convenient number to assess integer coupling by providing a single number for each gastric mill cycle. Integer coupling exists when the significand value is either close to zero or close to one. To compare the effect of temperature on integer coupling, we plotted the significand in a rose diagram (Fig. 7B). This representation is meaningful because rhythmic patterns are inherently comprised of repetitive circular patterns, and the end of one cycle represents the beginning of the next. We defined the beginning of the pyloric cycle as 0 and the end as 1. In this view, significand values near 0 and 1 coincide at the top of the diagram, providing a continuous measure for integer coupling. We plotted the significand of all gastric mill cycles and experiments separately for each temperature condition and measured the resulting vector strength. We found that, as long as the gastric mill rhythm was active, the resulting vector was significantly different from an equal distribution, supporting the Null hypothesis that most LG bursts occurred after an integer number of pyloric cycles (Fig. 7B). Thus, the two rhythms were integer coupled at all temperatures, even when only the STG was heated. However, we noted that the z values of our Rayleigh analysis diminished with higher temperatures, suggesting a wider distribution of significands and thus a weakened integer coupling. The weakened integer coupling was also evident from the wider spread of data points in the rose plots at 16 and 19°C (Fig. 7B).

Since integer coupling between the pyloric and gastric mill rhythms depends on LG disinhibition when the pyloric pacemakers inhibit Int1 *(Bartos et al., 1999)*, the LG burst should start at a fixed phase of the pyloric cycle. We tested whether this was the case by plotting the phase of the LG burst onset within the pyloric cycle, again using rose plots (Fig. 7C). A pyloric cycle was defined from the start of the PD neuron burst to the beginning of the subsequent PD burst (see Material and Methods). The cycle period was calculated using the average of the two cycles preceding the LG burst. The onset phase of the LG burst was then calculated as the time delay with which the LG burst began after the start of the PD burst, divided by the pyloric cycle period. As expected for integer coupling, the observed phases clustered within a small range at low temperatures. Like for the significands, the averaged resulting phase vector was statistically different from a random distribution at all temperatures (Fig. 7C). Thus, both measures, significand and phase, demonstrated that integer coupling was present. However, the range of observed phases increased at 16 and 19°C when compared to colder temperatures. While still statistically different from a random distribution and thus integer coupled, the coupling appeared to weaken at 16°C and above. Less stringent phase coupling between the rhythms could result from LG being less likely to produce action potentials as the rhythm slows and cycle periods become long (Figure 2). LG could thus be more prone to fail to properly initiate a burst, resulting in varying delays between the pyloric disinhibition and the LG burst onset.

Per definition, no integer coupling can exist between the pyloric and gastric mill rhythms at crash temperature when the gastric mill rhythm stops. However, since rhythms continue to function when STG and CoGs are heated together, or CabTRP Ia is present, we asked whether integer coupling is maintained when the rhythm is rescued. For analysis, we plotted the significand during temperature- and CabTRP Ia-mediated rescue, respectively. For the temperature rescue, we heated STG and CoGs together to crash temperature to rescue the rhythm (similar to Fig. 4A). For the CabTRP Ia rescue, we heated only the STG to crash temperature and then applied CabTRP Ia selectively to the STG to rescue the rhythm (similar to Fig. 6). For both rescue conditions, the significand vectors were statistically different from equal distribution (Fig. 7D), and z values were high (temperature rescue: 24.3, CabTRP Ia rescue: 47.2), indicating that integer coupling was present. We also found that the phase at which the LG started within the pyloric cycle was statistically different from a random distribution for both rescue conditions (Fig. 7E) with high z values (temperature rescue: 38.5, CabTRP Ia rescue: 48.9). Since neither heating the CoGs nor applying CabTRP Ia altered the coupling mechanism between the rhythms, our data further verifies that integer coupling is intrinsic to the STG circuits.

### Integer coupling becomes weakens with increasing temperature

We noted that the significands in the temperature rescue experiments, while significant, had a wider range than at most pre-crash temperatures and concurrently lower z-values in the Rayleigh analysis. This was surprising since we expected temperature rescued rhythms after warming STG and CoGs together to show a robust integer coupling, as *Powell et al. (2021)* suggested. To resolve this mystery, we looked at individual experiments rather than the averaged data. Figure 7F (left) shows original recordings of a temperature-rescued gastric mill rhythm at 19°C, and the simultaneously recorded pyloric pacemaker activity (PD). The colors indicate the timing of the LG burst within the pyloric cycle, i.e., its onset phase. Unexpectedly, LG’s onset phase was not constant, although the rhythm was regular and long-lasting. For example, the LG burst began at phase 0.65 for the first LG burst (top recordings), while the LG burst of the subsequent cycle (second recording from top) started at phase 0.18, suggesting that the two rhythms were not tightly coupled anymore. In contrast to this experiment, we also found examples where LG’s onset phase mainly remained constant in the rescued rhythm (Fig. 7F, right). Thus, there appeared to exist at least two data populations, one where strong integer coupling was maintained at higher temperature and another one where integer coupling was mostly absent despite regular rhythmic activity.

To exclude that the observed shift in phase-relationship in some preparations was not an idiosyncrasy of our specific approach where just parts of the preparation were heated, we compared our data to the publicly available data set from *Powell et al. (2021)*, where the whole preparation was heated uniformly. We re-plotted the significands from the Powell et al. data set in rose diagrams (Figure supplement S1A). In addition, we analyzed the phase relationship between PD and LG onset and calculated the phase vectors (Figure supplement S1B). To our surprise, we found very similar results to our experiments. At all temperatures, the significand and phase vectors in the Powell et al. data set were significantly different from random distributions, demonstrating that, on average, integer coupling was present. However, we also found that with increasing temperature, the range of significands and phases broadened.

Furthermore, when we looked at individual preparations, we found that some showed varying LG onset phases and increasingly so at higher temperatures, while others maintained a stable onset phase. Figure supplement S1C shows an example of one such experiment. The onset phase of the LG burst varied dramatically, from 0.23 (top recording) to 0.76 during the subsequent gastric mill cycle (second from top). Consequently, even when the whole nervous system is heated uniformly, two subpopulations of responses seem to be present. These results thus demonstrate that even in small circuits, idiosyncrasies exist between individuals, resulting in distinct responses to temperature challenges. This fits well with previously suggested variabilities between individuals concerning varying ion channel expression levels *(Goaillard et al., 2009; Schulz et al., 2006)* and responses to perturbations and physical variables such as pH and temperature *(Alonso and Marder, 2020; He et al., 2020)*, and the general idea of degenerative circuits *(Powell et al., 2021)*.

## Material and Methods

### Animals

Adult *Cancer borealis* were purchased from The Fresh Lobster Company (Gloucester, MA) and maintained in filtered, aerated artificial seawater (salt content ∼1.024g/cm^3^, Instant Ocean Sea Salt Mix, Blacksburg, VA) at 10 to 11°C with a 12-hour light-dark cycle. Animals were kept in tanks for at least 10 days prior to experiments.

### Solutions

*C. borealis* saline was composed of (in mM) 440 NaCl, 26 MgCl_2_, 13 CaCl_2_, 11 KCl, 11 Trisma base, and 5 maleic acid, pH 7.4–7.6 (Sigma Aldrich). In some experiments, 1 μM CabTRP Ia (GenScript, Piscataway, NJ) was added to the saline. Solutions were prepared from concentrated stock solutions immediately before the experiment. Stock solutions were stored at −20°C in small quantities. Measurements were taken after 45 min wash in/out.

### Dissection

Animals were anesthetized on ice for at least 30 minutes. The stomatogastric nervous system was dissected following standard procedures as described previously (Städele et al., 2015) and pinned out in a silicone-lined petri dish (Sylgard 184, Dow Corning). Preparations were continuously superfused with physiological saline (7-12mL/min).

### Temperature control

A petroleum jelly well was built around the STG to isolate it from the rest of the nervous system thermally (Fig. 1A). Temperature inside and outside the well was controlled independently with two saline superfusion lines, each heated by separate Peltier devices. Temperature was altered between 7 to max. 25°C in 3°C increments and changed by ∼1°C/min ± 0.2°C. The preparations were allowed to acclimate to the new temperature for 5 minutes before gastric mill rhythms were elicited between each temperature step. Temperature was continuously measured close to the STG and CoGs with separate temperature probes (Volt-craft 300K, Conrad Electronik, Germany). Saline inflow to the nervous system was positioned within 1 cm of the STG so that the measured temperature at the point of inflow was approximately that of the ganglion somata. Three different temperature alternation experiments were performed. 1.) Temperature in the STG well was altered, while the surrounding nervous system was kept constantly at 9°C. 2.) Temperature of the surrounding nervous system was changed (=bath), while the STG circuits were continuously kept at 10°C. 3.) Temperature of both the STG circuits and the surrounding nervous system was altered simultaneously.

### Electrophysiological recordings

Extracellular recordings were performed from nerves via stainless steel pin electrodes procedures (Städele et al., 2017). Stretches of the *lgn, dgn, mvn, pdn*, and ions were electrically isolated from the bath with Vaseline wells. Voltage signals were recorded, amplified, and filtered using an AM Systems amplifier (Model 1700; Everett, WA) and digitized at 10 kHz with a Power 1401 (CED, Cambridge, UK).

### Gastric mill rhythm stimulation

To elicit a gastric mill rhythm, we bilaterally stimulated the dorsal posterior esophageal nerves (*dpons*). Extracellular stimulation was performed using standard procedures (Städele et al., 2017) using a Master-8 stimulator (AMPI, Jerusalem, Israel) with 10 stimulus trains of six seconds duration at 0.6 Hz. Intratrain stimulation frequency was 15 Hz and each stimulus pulse was 1ms in duration (Beenhakker et al., 2004). The stimulation amplitude was set to 1.5 times the minimum voltage needed to elicit a response of STG neurons. The stimulus site was maintained at constant temperature throughout the whole experiment.

### Data analysis, statistical analysis

Electrophysiological files were recorded, saved, and analyzed using Spike2 (Version 7.18, CED, Cambridge, UK), and original Spike2 and Matlab scripts. No spike sorting was necessary as all neurons could be individually identified on the respective nerve recordings, or from intracellular recordings. In case of MCN1 recordings from the *ion*, the large action potentials of the esophageal motor neuron were removed prior to measuring MCN1 following established protocols (Hedrich et al., 2011). Bursts in all neurons were defined using inter-spike intervals and spike numbers. Inter-spike intervals longer than 0.25 seconds were defined as a pause between bursts (inter-burst interval) for PD. Inter-spike intervals longer than 1 second were defined as a pause between the bursts of LG and DG. For all neurons, a minimum of 2 spikes was required to define a burst. Cycle periods were calculated from the beginning of one burst to the beginning of the next (PD for pyloric cycle period, LG for gastric mill cycle period). Duty cycles were calculated by dividing the burst duration by cycle period.

All data was normally distributed, and we employed parametric tests (ANOVA, student’s t-test, Rayleigh z test). In cases where different variables were measured in individual animals, repeated measures tests were considered. The factors and specific designs of each ANOVA are described in the corresponding figure legends. Statistical tests were performed using SigmaStat (version 11; Systat Software, San Jose, CA). For ANOVA, statistical tests are reported in the format: statistical test, F(degrees of freedom, residual)=F value, p value, post hoc test, number of experiments. “N” denotes the number of preparations, while “n” denotes the number of trials. Post hoc tests are at a significance level of 0.05. Data was prepared in Excel and MATLAB and finalized in Adobe Illustrator (version 25.4.1; Adobe Inc.). In figures, data is presented as mean±SD unless otherwise noted. In some figures, average data are connected by a line when recorded in multiple conditions to indicate paired data.

### Integer coupling analysis

To determine whether integer coupling was present or not, we calculated the significand. A detailed description is given in Powell et al (2021). In short, the significand is calculated by dividing the gastric mill cycle period by the pyloric cycle period, and then taking the decimal part of the result. When zero or one, integer coupling is present because an integer number of pyloric cycles is present in one gastric mill cycle.

As a second measure for integer coupling, we used the phase at which the LG bursts started within the pyloric cycle. Phase was calculated by first measuring the time delay at which the LG burst started after the last preceding PD burst. This time was then divided by the pyloric cycle period, resulting in a number between 0 and 1 (the onset phase of LG). In any given condition (e.g., at one temperature), LG onset phase must remain the same for each burst if integer coupling is present.

To determine if any set of significands or LG onset phases was significantly different from a uniform distribution, we created rose (circular) plots and carried out a Rayleigh z test (Ruxton, 2017). Significands and phases were plotted in a circle from 0 to 1, and mean directional vectors were calculated. Uniform distribution indicates the absence of integer coupling, a rejection of the null hypothesis rejects uniformity and supports integer coupling.

## Additional Information

### Author contributions

Carola Städele: Conceptualization, Data curation, Software, Investigation, Visualization, Methodology, Writing - original draft, Writing - review and editing. Wolfgang Stein: Conceptualization, Visualization, Supervision, Funding acquisition, Validation, Project administration, Writing - original draft, Writing - review and editing.

### Additional files

pdf file containing Figure supplement S1.

