## Supplemental Figure S1 for "Neuromodulation enables temperature robustness and coupling between fast and slow oscillator circuits in *Cancer borealis*"


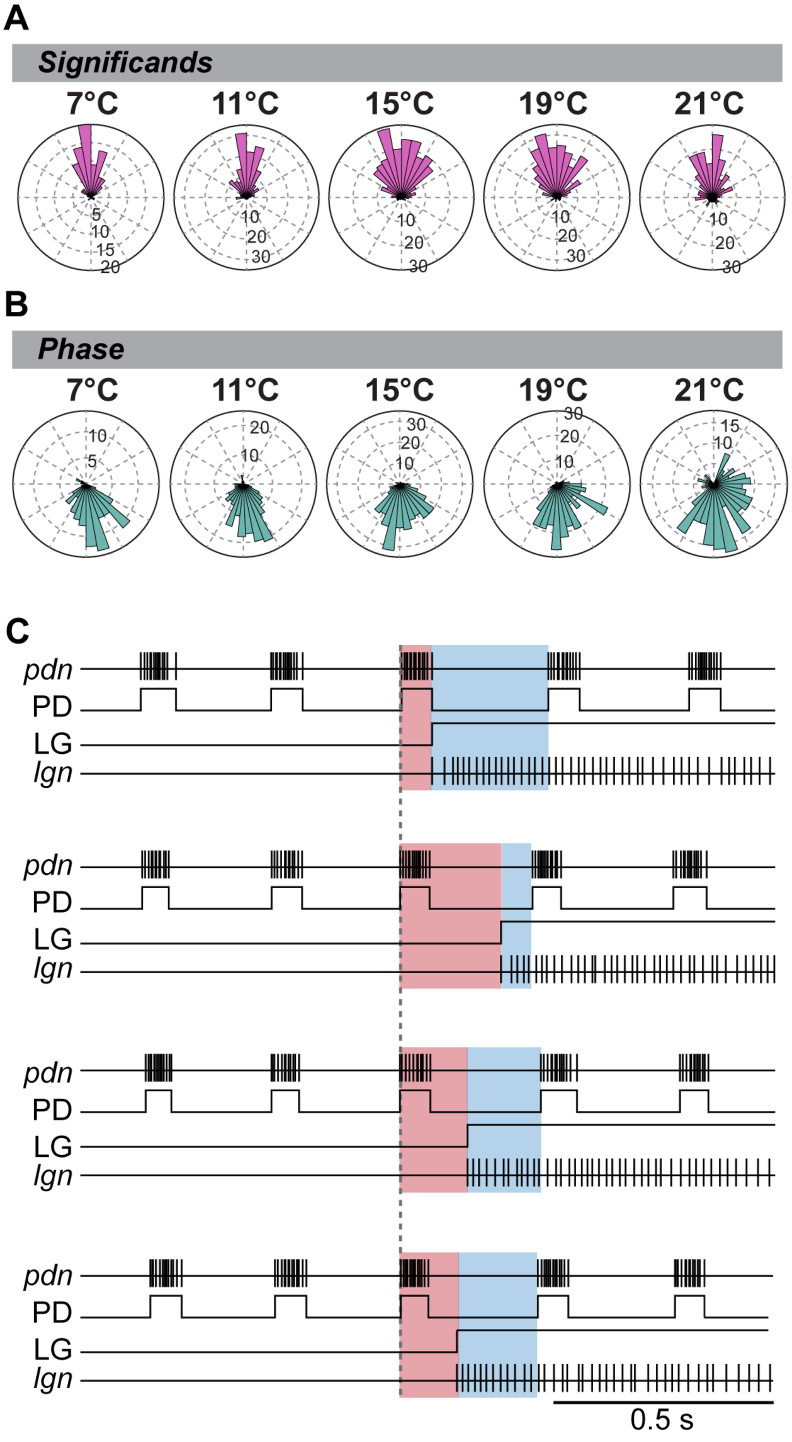


**Figure supplement S1: Data from Powell et al. (2021), reanalyzed. (A)** Significands of all gastric mill cycles and animals at various temperatures. Rayleigh z Test: 7°C: Rayleigh $z_{70.5}$, $p<0.001$, 11°C: Rayleigh $z_{108.5}$, $p<0.001$, 15°C: Rayleigh $z_{135.3}$, $p<0.001$, 19°C: Rayleigh $z_{146.8}$, $p<0.001$, 21°C: Rayleigh $z_{66.3}$, $p<0.001$. **(B)** Phase of the LG burst start within the pyloric cycle at various temperatures. All gastric mill cycles and animals are shown. Rayleigh z Test: 7°C: Rayleigh $z_{60.8}$, $p<0.001$, 11°C: Rayleigh $z_{112.8}$, $p<0.001$, 15°C: Rayleigh $z_{153.6}$, $p<0.001$, 19°C: Rayleigh $z_{147.5}$, $p<0.001$, 21°C: Rayleigh $z_{53.3}$, $p<0.001$. **(C)** Example recordings for one animal at 21°C. The LG burst is shown with a recording of the *lgn* and schematized LG bursts (LG). Four individual LG burst onsets are displayed (top to bottom). Pyloric cycles are shown by recordings of the PD neurons (*pdn*) and schematized PD bursts (PD). The dashed vertical line indicates the beginning of the last PD burst before the LG burst onset. The time delay between PD onset and LG onset is highlighted in red. The remainder of the pyloric cycle is marked in blue. A change in the ratio of red and blue illustrates a shift in the phase relationship between PD and LG onset. Phases varied dramatically.


*Literature cited:*

**Powell D, Haddad SA, Gorur-Shandilya S, Marder E**. 2021. Coupling between fast and slow oscillator circuits in *Cancer borealis* is temperature-compensated. *Elife* **10**:1–23.
